# MSIGNET: a Metropolis sampling-based method for global optimal significant network identification

**DOI:** 10.1101/260844

**Authors:** Xi Chen, Jianhua Xuan

## Abstract

In this paper, we propose a novel approach namely MSIGNET to identify subnetworks with significantly expressed genes by integrating context specific gene expression and protein-protein interaction (PPI) data. Specifically, we integrate differential expression of each gene and mutual information of gene pairs in a Bayesian framework and use Metropolis sampling to identify functional interactions. During the sampling process, a conditional probability is calculated given a randomly selected gene to control the network state transition. Our method provides global statistics of all genes and their interactions, and finally achieves a global optimal sub-network. We apply MSIGNET to simulated data and have demonstrated its superior performance over comparable network identification tools. Using a validated Parkinson data set we show that the network identified using MSIGNET is consistent to previously reported results but provides more biology meaningful interpretation of Parkinson’s disease. Finally, to study networks related to ovarian cancer recurrence, we investigate two patient data sets. Identified networks from independent data sets show functional consistence. And those common genes and interactions are well supported by current biological knowledge.

## Introduction

Large scale protein-protein interaction (PPI) networks of Homo species including nearly 9,000 genes with 39,000 interactions are now available through yeast two-hybrid studies (1). These interactions advance our interpretation about the response of protein complexes (2), signal transductions (3,4) and signalling pathways (5) to extracellular impact caused by certain disease. In recent years, cancer related network analysis promotes the importance of context specific network identification (6–9). Even though many of genes have been individually studied through biological experiments, interactions between them give us more information about the impact of signalling pathway on the cancer development. Due to the growing size of public tumour samples, i.e. TCGA database, at the gene expression level, it becomes feasible to study the difference between normal samples and disease samples. Moreover, for drug resistance or cancer recurrence, besides genomic analysis focusing on somatic mutations (10,11), rewiring of gene interactions in recurrent vs. non-recurrent may guide us to identify recurrence related signalling pathways. And network based prediction has been demonstrated to be more power than gene based classification (12).

Identifying differentially expressed gene network is long lasting topic in system biology field. A few methods published in recent years combine both kinds of scoring together to reflect the differential expression from network aspect. RegMOD (13) uses signal-noise ratio to for node score and correlation coefficient for edge score. Based on a regression model, it calculates each gene’s active score function based on the diffusion kernel. PNA (14) utilize smultivariate logistic functions to form a weight matrix containing activity information for both nodes and edges, and then, generate principle subnetworks with orthogonal non-negative matrix factorization. COSINE (15) employs F-statistics based approach to jointly measure the condition specific changes of both nodes and edges. Based on available methods, z-score in (16) is the most common way to construct the node score. And for edge, both correlation coefficient and mutual information make sense. Identifying optimal sub-network is proved to be a NP hard problem (6). With the network score design, a searching process is needed for network identification. A global statistical method to evaluate the confidence of each node and edge in current network is what we need to investigate the network identification problem. Metropolis sampling/ Monto Carlo Markov Chain is a better choice to all above problems.

In this paper, we propose a Metropolis sampling based SIGnificant NETwork (MSIGNET) identification method. The network score involves node significance, edge correlation as well as mutual information. We specifically design a conditional probability function to make the Markov Chain have a high probability to transit to a more meaningful state (a higher network score). Through the simulation study, we show that MSIGNET did a better job on subnetwork identification, where several isolated local optimal modules co-exist. We further apply MSIGNET to real data, including one Parkinson patient data set and another two ovarian cancer patient data sets. In the Parkinson data set, MSIGNET not only reproduce the network identified using another method, it provides more unreported but biological meaningful interactive genes associated with Parkinson disease. The similarity between MSIGNET networks of two ovarian cancer data sets demonstrates the robustness of the proposed method. The common network including many known ovarian cancer genes provides insights to the hidden module driving ovarian cancer development.

## Methods

### Metropolis sampling based significant network identification

By integrating both gene expression data and PPI network, we calcualte a network score consisting of three components as node z-score, a linear edge score (spearman correlation), and a non-linear edge score (mutual information). The non-linear score a a higher order portray of the similarity between connected genes. Then, we use a Metropolis sampling based searching method. Instead of using random walking for node or edge selection, we propose a conditional probability density function. This function can greatly shorten the burning time, a big concern for most sampling-based approaches. Finally, we utilize bootstrapping to combat the outlier impacts and significant test to lower the randomness of our identified network. To validate the performance of MSIGNET, we first simulated gene expression data with different quality and three networks with different topologies to test MSIGNET performance on both node and edge identification. Then, we applied MSIGNET to a Parkinson gene expression data set (17) with validated gene interactions. Finally, we applied MSIGNET to TCGA (https://cancergenome.nih.gov/) ovarian cancer patient gene expression data and compared the identified network to the network identified from another ovarian cancer data set with a similar treatment (18).

### Network Score Design

MSIGNET is developed to identify significant network with genes significantly over or lower expressed in a certain condition. Therefore, there should be two groups in a gene expression data for differential analysis. Given a gene expression data set {*X_n,s_ | n = 1 ~ N, s = 1 ~ S*} including *N* genes and *S* samples, the s-th sample can be classified to one of two groups according to phenotypes or treatment, with label *c(s)* = 0 or *c(s)* = 1. Then, we use the two-sample t-statistics to calculate a *p _ value(n)* for *n*-th node and further transfer this *p _ value(n)* into a z-score *z(n)* using an inverse Gaussian function Φ^−1^ as follows:

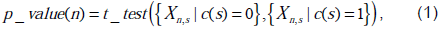

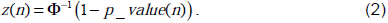

For the edge score, we calculate the Spearman rank correlation coefficient *ρ*(*n,m*) to portray the relationship of two connected genes, i.e. n-th gene and m-th gene. Then, we converted the correlation coefficient to a t-score in order to assess its significance.

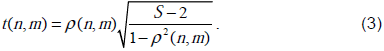

*t* score follows a student t distribution with a degree of *s − 2*. It significance p-value is calculated as *p _ value(n,m)*. Similarly, using an inverse Gaussian function Φ^−1^ we transfer this *p _ value(n,m)* into an edge score *e(n, m)*.

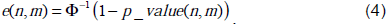

Assuming that the whole PPI network has *N* genes (nodes) and *L* edges, for a sub-network *i* with *N_i_* nodes and *L_i_* edges, we first calculate a network node score *Z_N_i__* and edge score *E_L_i__*.

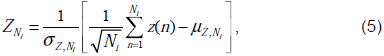

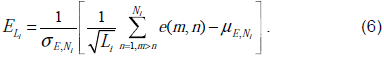

However, the edge score is still based on the linear relationship of two genes’ expression. Mutual information can provide a higher order portray of the dependence of pairwise genes. The analysis of information flow using mutual information of edges (19) is more advanced than the linear correlation-based edge analysis. The information of a network is calculated by adding the mutual information of all edges together as follows:

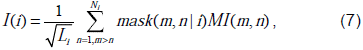

where *MI(m,n)* is the estimated mutual information based on the expression data of genes *n* and *m* using a method from (20).

Finally, a joint probability including node score, edge score as well as the information contained in the network is defined as follows:

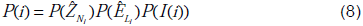

### Metropolis Sampling based Network Searching

In this paper, we design a Bayesian framework and use Metropolis sampling to identify a global optimal network by integrating context specific gene expression with the human PPI network. The sampling process can be modelled as a Markov process. We define a node vector *V* with length of *N* and an edge matrix *W*. The PPI network is quite sparse so we assume there are *M* interactions in the matrix *W*. To initiate the Markov chain, we randomly select a small number of connected nodes, i.e. 30. By assuming the network in the *i*-th round of sampling has *V_i_(n) = 1* if node *n* is sampled, between two consecutive rounds of sampling, the network jumps from state to state *j*, by changing only one node and one edge. It is possible that changing a node will cause multiple changes of edges. In this framework, we only allow a lead node of the current network deleted (the delete of a leaf node does not break the network). In each round of sampling, we can either add a node or delete a node with a prior probability 0.5. The ‘jump’ of Markov chain from state to state *j* is controlled by a conditional probability *P*(*j|i*). *P*(*j*| *i*) is equavalent to *P*(*m*| *i*) since there is only one egde (*m*-th edge) changed. In the following part, we design the likelihood function *P*(*m*| *i*) for adding and deleting, respectively.

For node addition, first, we select a node *n* in current network as a seed (dark red in Fig. 1(a)). The prior probability for this selection is uniform, which means all the nodes in the current network are equally important. Then, all neighbour nodes to the seed excluding those nodes already in current network are treated as candidates, with a total number of *N_Link_*, which is 4 in Fig. 1(a). The score of a candidate neighbour node and its dependency (edge score) to the seed node jointly reflect the contribution to the current network. Therefore, a joint probability of two scores (each follows a Gaussian distribution) *P*(*z*(*m*)) *P*(*e*(*n, m*)) is used to compare all candidates. Given seed node *n* and current network state *i*, we define a proposal function as follows:

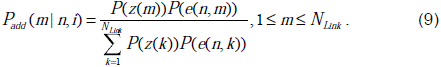

**Figure 1.**
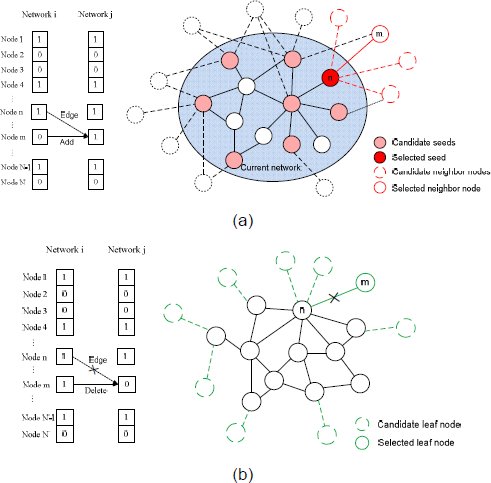
Node adding and deleting during the sampling process: (a) Add a node to current network according to the ‘adding’ conditional probability density function; (b) Delete a node from current network according to the ‘deleting’ conditional probability density function.

According to Eq. (9), we propose a new node as well as its edge based on the current network, as the solid red line in Fig. 1(a).

For deleting a node, different from the seed node selection for node addition, all leaf nodes in the current network become candidates. Considering that the size of current network is small (i.e. 30) and the network is quite sparse, there should be a number of leaf nodes to select. As shown in Fig. 1(b), we intend to delete the leaf node that contributes the least to current network. For each leaf node *m*ϵ[1, *N_leaf_*], we calculate a joint probability [1 - *P*(*z*(*m*))][1 - *P*(*e*(*m,n*))], where *n* is the connected node for leaf node *m*. If this probability is high, this leaf node is neither expressed differently nor intensely related to its connected node in current network. Therefore, the proposal functional for deleting node or edge is defined as follows:

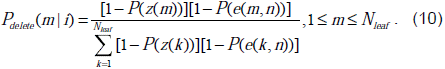

Based on Eq. (9) or (10), we propose a node (and an edge) for adding or deleting, however whether this proposal is accepted depends on the overall performance of a network. We define the following criterion to determine whether we accept the proposal:

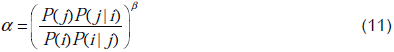

If *α* ≥ 1, we will accept the proposal; otherwise, we accept the proposal by a probability *α*. The parameter *β* controls the acceptation rate. A high value of *β* will lower the probability for adding a ‘negative’ node or deleting a ‘positive’ node. After each round of sampling, node vector *V* and edge matrix *W* will be updated. After a long sampling process, *V* and *W* would provide related but different statistics on nodes and edges in a network.

### Bootstrapping

For real data analysis, especially for cancer related gene expression data set, the impact of limited number of patient samples and data noise cannot be ignored. The gene expression data is quite diverse due to the characteristics of each patient. Any outliers or errors existing in the gene expression data will impact our computational results. To address statistical confidence for network identification, we use bootstrapping technique to run MSIGNET 100 times by bootstrapping gene expression samples. The mean estimate of 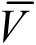 and 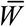 across all 100 replicates are used as the final statistics on nodes and edges.

## Results

### Gene Network and Expression Data Simulation

To simulate different scenarios of protein-protein interactions in real networks, we construct underlying ground truth networks by bridging several gene modules observed in the Human Protein Reference Database (HPRD Release 9, http://www.hprd.org/download). We want to test whether MSIGNET and competing methods can identify a “global optimal” network with two or more significant modules. We design three networks with different topologies.

Network 1 is a small network with two close DE modules bridged by one EE module. There are 125 ‘true’ nodes in two DE modules and 62 ‘false’ nodes in the EE module. A ‘true’ edge is defined as an edge connecting two DE nodes. In total there are 360 true edges out of 1010 edges in Network 1 with only 9 nodes shared by the two DE modules.

Network 2 also has two close DE modules but surrounded by several other EE gene modules with a higher density. There are 127 DE nodes and 927 true edges in the network with totally 260 nodes and 1663 edges. Compared to Network 1, the number of DE genes is similar, but the edge density is increased. Also, the number of background EE genes is increased with more modules between or surround the two DE modules. The overlap between two DE modules are just 8 nodes.

Network 3 has two distant DE modules separated by a group of EE modules. There is very little overlap between two DE groups. In total we have 220 significant nodes and 1513 true edges in the network (2518 nodes and 15266 edges). This is quite a challenging case to test whether the method could avoid local optimal during the network searching. If a method suffers from local optima issue, it will stuck into one module and cannot jump to the other DE module through several paths with EE genes only.

We simulated genes states as differentially expressed (DE) or equally expressed (EE) with a Markov random field (MRF) model (21). Here, an advantage of using MRF model is that we can control the consistency between simulated DE genes and ground truth DE genes by adjusting the weight function in MRF model. Then, using a Gamma-Gamma (GG) model (22), we generated gene expression data based on the states of genes with additive Gaussian noise. Here, we set the signal-to-noise ratio (SNR) as 4dB and the weight of MRF-GG model as 10. After 1e5 rounds of sampling, we get the samples at all nodes and edges, respectively. ROC curves of MSIGNET are shown in Fig. 2 for node and edge evaluations, respectively, mainly in comparison with RegMOD. For the remaining competing methods, since they directly output a final network, a F-measure (2*precision*recall/(precision+recall) of each method on ‘true’ node or edge identification is summarized in Table 1.

**Figure 2.**
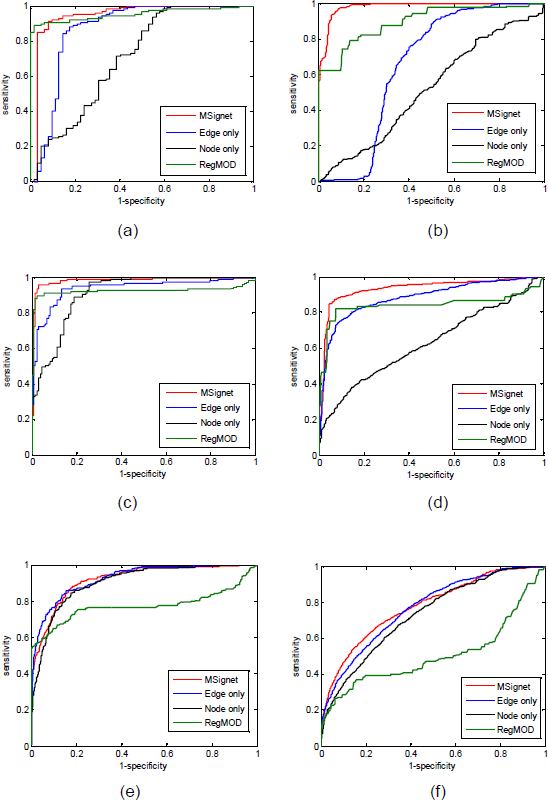
ROC curves of node and edge identification. (a) Network 1 node identification performance; (b) Network 1 edge identification performance; (c) Network 2 node identification performance; (d) Network 2 edge identification performance; (e) Network 3 node identification performance; (f) Network 3 edge identification performance.

**Table 1.** F-measure of competing methods for node and edge identification.

| Methods |  | **MSGINET** | COSINE | MATISSE | Heinz | jActive | GiGA | DeGAS |
| --- | --- | --- | --- | --- | --- | --- | --- | --- |
| Network 1 | Node | **0.952** | 0.524 | 0.920 | 0.910 | 0.972 | 0.840 | 0.874 |
|  | Edge | **0.819** | 0.336 | 0.538 | 0.718 | 0.812 | 0.657 | 0.715 |
| Network 2 | Node | **0.961** | 0.493 | 0.918 | 0.922 | 0.938 | 0.940 | 0.700 |
|  | Edge | **0.896** | 0.314 | 0.543 | 0.895 | 0.891 | 0.894 | 0.509 |
| Network 3 | Node | **0.877** | 0.153 | 0.667 | 0.665 | 0.491 | 0.420 | 0.562 |

For Network 1, although in Fig. 3 (a) for node identification, MSignet and RegMOD have similar ROC performance, as can be found from Fig. 3 (b) that MSIGNET is much better than RegMOD on edge identification. In Table 1, only jActive is comparable to the proposed MSIGNET. Using Network 1, we show that MSIGNET provides comparable performance on node identification as most other methods, and also it provides a more accurate selection on network edges.

**Figure 3.**
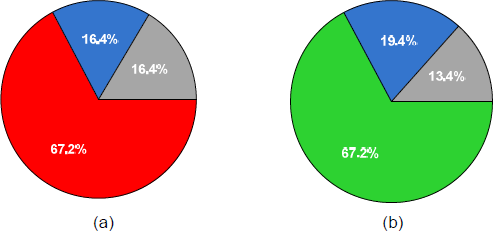
Parkinson’s disease network validation distribution: (a) up regulated genes; (b) down regulated. ‘Red’ pie and ‘Green’ pie are the percentage of validated up-regulated and down-regulated genes, respectively; ‘Blue’ pie and ‘Grey’ pie are missed out gene by our method and original network coverage, respectively.

**Figure 4.**
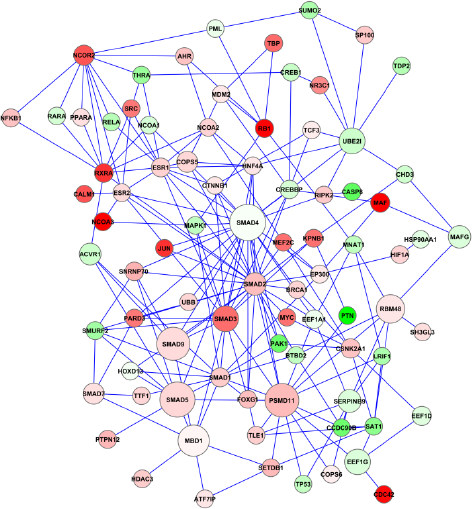
Common network from both TCGA and Duke data sets. Node colour and size are based on the results of the TCGA gene expression data set. Red genes are overexpressed in early recurrence group while green ones are overexpressed in late recurrence group. The size of each node and the width of each edge respectively represent the sampling frequency of MSIGNET.

It can be seen from Fig. 3 (c) and (d) that for Network 2, MSIGNET is more advanced than RegMOD on both node and edge identification. Furthermore, as shown in Table 1, methods like COSINE and DeGAS are actually stuck in one module only with a recall performance around 0.5. Methods Heinz, jActive and GIGA work well in this network, but MSIGNET still provides a better performance.

Network 3 is very challenging since to achieve global optima, the method needs to go through the EE module frequently. Some ‘bridge’ edges have to be sampled to link the two separated DE modules. Obviously, in this case the advantage of using Metropolis sampling is maximized. The ROC performance of MSIGNET is significantly higher than RegMOD, as shown in Fig. 3(e) and Fig.3(f). RegMOD did a bad job on the edge identification, which is mainly due to the isolated distribution of significant nodes in Network 3. In Table 1, COSINE doesn’t converge with genetic algorithm, which provides a large network containing only 10% significant nodes; MATISSE, Heinz and DeGAS also stuck in one DE module with a quite low recall performance. jActive and GiGA both encounter the local optima issue and further, their precision even in one DE module is low. Through Networks 1 to 3, we demonstrate that MSIGNET provides a better solution to the global optima network identification.

### Validating Networks Identified from a Parkinson Gene Expression Data Set

In (17), Parkinson is investigated by applying DeGAS to gene expression data to identify disease associated networks, one network including 73 genes over-expressed genes in disease samples and another network including 67 depressed genes only. We downloaded their gene expression data from (23) with 18 normal samples and 29 disease samples. Based on the fold change of each gene between normal group and disease group, there are 5551 genes in the original PPI network showing up-regulation and 2712 genes in the PPI network showing down regulation. The network scale is quite large. We looked into details about the distribution of these genes identified in [17] in the PPI network. For the 73 up-regulated genes, most of them are falling within two jump neighbour genes around YWHAB; while for the 67 down-regulated genes, they are falling within two jump neighbour genes around HD. Therefore, we prune the original PPI by cutting a subnetwork including 1059 two jumps neighbour genes around YWHAB and another subnetwork including 1538 genes around HD. There are 61 up-regulated and 58 down-regulated genes validated in (17). We applied MSIGNET to each network and sort all genes based on sampling frequency. We select the top 150 genes in each group and compare to the results in (17). There are 49 validated up-regulated genes, as shown in Fig. 3 (a), ‘Red’ pie, and 45 validated down-regulated genes as shown in Fig. 3 (b), ‘Green’ pie. The ‘Blue’ pie represents unsuccessfully validated genes by our method while the ‘Grey’ pie is missed out due to limited network information. In Fig.3 (a), 80% of the up-regulated genes validated by [17] are successfully captured by MSIGNET, with a Fisher test p-value 5.6e-34. Similarly, in Fig.4 (b), 77% of down-regulated genes are finally identified by MSIGNET, with a p-value 4.9e-37.

Further, for the top 150 up-regulated genes reported by MSIGNET, we did functional enrichment analysis using David (https://david.ncifcrf.gov/). Cancer pathway, cell cycle and MAPK signaling pathway are significantly enriched. In addition, there are many genes evolving into cellular processes as cell cycle, cell growth and cell proliferation. And RNA splicing, regulation of apoptosis, cellular response to stress and regulation of cellular protein metabolic process are also enriched. Functionally, our results are consistent with (17). In the down-regulated network, for the top 150 genes reported by MSIGNET, here are more Parkinson’s disease related genes, totally 11 gene with p-value 3.7e-5. While in (17), only 6 genes were reported with p-value 2.4e-3. In addition, both Alzheimer’s disease (14 genes with p-value 1.5e-6) and

Huntington’s disease (16 genes with p-value 1.2e-7) are enriched. They are closely related to Parkinson’s disease and usually happen to people over 60 years old. Through functional annotation, the 159 genes also evolve in neurological system regulation (20), leaning, memory and behavior (10), cell cycle (14) and cell death or apoptosis (20). All these functions are closely related to the development of Parkinson’s disease.

### Network Identification using Ovarian Cancer Gene Expression Data Sets

We further investigated the significant network identification problem in cancer research. It is well known that currently, women suffer from high risk from ovarian cancer, which is very aggressive and usually can only be treated by chemotherapy. But the recurrence usually comes after 2 or 3 years. Current treatment is not effective enough to lower the risk of recurrence. The mechanism controlling the development of ovarian cancer is not clearly understood. Thence, it is necessary to focus on the genes which are differently expressed between early recurrence and late recurrence.

We first downloaded a ovarian cancer gene expression data set from the TCGA website. Totally there are 589 patient samples. We selected the samples with chemotherapy and based on the survival information, we further grouped those samples into early recurrence group (72 samples with survival time < 2 years) and late recurrence group (131 samples with survival time > 4 years). TP53 is proved to play an important role in ovarian cancer. Therefore, we selected all genes within two jumps from TP53 in the original PPI and after integrating with gene expression data, we finally obtained 2240 candidate genes and 11798 edges between them. We apply MSIGNET to this data set. Since the real patient data is quite noisy, to lower the impact of outliers in either group, we conducted 100 rounds of bootstrapping. We selected nodes with the 10% average sampling frequency after bootstrapping. To ensure connectivity, top 25% edges are filtered based on average sampling frequency after bootstrapping. TCGA data is generated by several groups so it is quite noisy, and the batch effect may also exist. To improvement the robustness of our network results, we further applied MSIGNET to another ovarian cancer data set GSE3149 (http://www.ncbi.nlm.nih.gov/geo/). The grouping of patients is similar to what we have done for the TCGA data set. We got 33 samples in early recurrence group and 56 samples in late recurrence group. We applied MSIGNET to this data set and generated another network. Again, top10% nodes and 25% edges were selected.

The common network of two studies is shown as Fig.4. In this network, SMAD family, NCOR2, NFKB1, UBE2I, TP53, MYC, BRCA1, ESR1, ESR2, JUN, MAPK1 and NR3C1 are reported to be related to ovarian cancer in different publications. An important gene associated with ovarian cancer is BRCA1, and among its related pathways are NF-KappaB Family Pathway and TGF-beta signaling pathway. For the upper part, NCOR2 serves as a hub closely connected with several significant genes. Aberrant expression of NCOR2 is associated with certain cancers. NFKB1, a very important ovarian cancer related gene, is shown as a small bub in the network (upper left corner), too, which is overexpressed in early group. Even though the fold change of itself is not quite significant, it has a direct connection to RXRA, which is proved to be overexpressed in most ovarian cancer patients. SPTBN1 is significantly overexpressed in early group and closely connected with CSNK2A1. CSNK2A1 has been found to be increased at protein level and up-regulated at the level of enzyme activity in the majority of cancers. And it is directly linked with BRCA1. Doing functional annotation using David we identified 13 genes enriched in TGF-beta signaling pathway with p-value 1.6e-11. It is not strange to find cell cycle (p-value 2.84e-8), cell death (p-value 7.61e-6) and apoptosis (p-value 2.06e-10) due to their importance of cell development. Several cancer related pathways are highlighted and in total 21 genes are enriched in the pathway of cancer. It is interesting that most of these cancer related genes also get involved in the regulation of cell proliferation (14 genes) and cell growth (6 genes). And MAPK signaling pathway is enriched with p-value 3.58e-3.

## Discussion

We introduce a Metropolis sampling-based method to identify a global optimal significant network by integrating gene expression data and PPI network. The superiority of MSIGNET is mainly reflected from the following aspects: (1) we incorporate the mutual information between genes into the network score. This builds a higher order description of node dependency besides the linear correlation of gene expression; (2) we use a mCmC process in the network searching. Compared to simulated annealing or greedy search in our simulation study, this approach did a better job when there are multiple local optimal modules in the network; (3) instead of using a random walk model, MSIGNET builds a conditional probability function for adding or deleting nodes or edges. This design significantly lowers the randomness of node searching and speeds up the convergence the Markov chain. However, a notable issue existing in MSIGNET is the time efficiency. First of all, the estimation of mutual information costs some time, much slower than the linear correlation. Second, due to the use of Metropolis sampling, a long burning process is necessary to get a converged Markov chain. Therefore, to make MSIGNET work efficiently, users are required to control the network size to one or two thousand genes. But if the computing time is not a big concern in order to get global optimal results, especially in cancer data analysis, MSIGNET can be applied to any network without size limitation.

